# Controlled Noise: Evidence of Epigenetic Regulation of Single-Cell Expression Variability

**DOI:** 10.1101/2024.04.10.588957

**Authors:** Yan Zhong, Siwei Cui, Yongjian Yang, James J. Cai

## Abstract

**Motivation:** Understanding single-cell expression variability (scEV) or gene expression noise among cells of the same type and state is crucial for delineating population-level cellular function. While epigenetic mechanisms are widely implicated in gene expression regulation, a definitive link between chromatin accessibility and scEV remains elusive. Advances in single-cell techniques now enable simultaneous measurement of scATAC-seq and scRNA-seq within individual cells, presenting an unprecedented opportunity to address this gap.

**Results:** This paper introduces an innovative testing pipeline to investigate the association between chromatin accessibility and scEV. The pipeline hinges on comparing the prediction performance of scATAC-seq data on gene expression levels between highly variable genes (HVGs) and non-highly variable genes (non-HVGs). Applying this pipeline to paired scATAC-seq and scRNA-seq data from human hematopoietic stem and progenitor cells, we observed a significantly superior prediction performance of scATAC-seq data for HVGs compared to non-HVGs. Notably, there was substantial overlap between well-predicted genes and HVGs. The gene pathways enriched from well-predicted genes are highly pertinent to cell type-specific functions. Our findings support the notion that scEV largely stems from cell-to-cell variability in chromatin accessibility, providing compelling evidence for the epigenetic regulation of scEV and offering promising avenues for investigating gene regulation mechanisms at the single-cell level.

**Availability and implementation:** The source code and data used in this paper can be found at https://github.com/SiweiCui/EpigeneticControlOfSingle-CellExpressionVariability.

## 1. Introduction

Single-cell RNA sequencing (scRNA-seq) has become a crucial and powerful tool for characterizing gene expression at the individual cell level, offering unprecedented opportunities to study gene regulation. While many studies have focused on examining the mean expression levels of genes, researchers have increasingly recognized the importance of cell-to-cell variability in gene expression, also known as single-cell expression variability (scEV) or gene expression noise, among cells of the same type and state [1-3]. Dueck et al. [4] proposed the “variation is function” hypothesis, suggesting that scEV plays a significant role in manifesting population-level cellular function. Their hypothesis highlights that scEV pertains to the diversity within a highly homogeneous population of cells rather than to the diversity of a mixed cell population with different, clearly distinct cell types that have already been recognized in previous studies. Osorio et al. [5] validated the “variation is function” hypothesis by identifying highly variable genes from homogeneous cells of different cell types and demonstrated that, for each cell type, the functions of highly variable genes are enriched with biological processes and molecular functions precisely relevant to the biology of the corresponding cell type. Other studies also support the functional importance of scEV. For instance, Wiggins et al. [6] reported an increase in scEV in BRCA1-associated breast tumors, providing new insights into the associations between genes and disease phenotypes.

Recently, with advancements in single-cell sequencing technology, researchers have been able to simultaneously assay both chromatin accessibility and transcriptomic profiles within individual cells using scATAC-seq and scRNA-seq, respectively. The scATAC-seq data provide valuable complementary information to the scRNA-seq data and enable a more intricate examination of the relationship between epigenetic modifications and gene expression dynamics at the single-cell level [7-8]. Granja et al. [9] developed a gene score based on the aggregation of scATAC-seq peaks that overlap with the gene window on open chromatin, which serves as a measure of gene expression control. They discovered a significant overall Pearson correlation between the constructed gene scores and gene expression levels. Mitra et al. [10] proposed a regularized Poisson regression method to predict scRNA-seq data using scATAC-seq data. Their approach showed improved prediction performance on both high- and low-coverage multiome datasets compared to gene score methods. More recently, Saelens et al. [11] employed neural network architectures to predict scRNA-seq data and identified differentially accessible chromatin regions between different cell types.

The coassay of scATAC-seq and scRNA-seq within the same cells offers an opportunity to study the epigenetic regulation of scEV. However, it is still unclear whether chromatin accessibility, as measured by scATAC-seq, impacts scEV. If so, how? There are several challenges in studying this topic. First, measuring the effects of scATAC-seq on scEV is difficult since the relationship between chromatin accessibility and gene expression is still an active research topic. A new measure needs to be proposed to solve this problem. Second, the source of scEV comes from both genetic and nongenetic factors and from both random and nonrandom effects [12]. Eling et al. [13] noted that accurately defining, measuring, and disentangling the stochastic and deterministic components of cell-to-cell variability is challenging. Third, scATAC-seq data is typically extremely sparse and high-dimensional with large variation [14]. Thus, further robust analysis is needed to explore the relationship between scATAC-seq and scEV.

In this paper, we propose a novel pipeline to investigate the notion of “the epigenetic regulation of scEV”. Our key idea is to use chromatin accessibility peaks (scATAC-seq data) to predict gene expression (scRNA-seq data) in single cells and compare the prediction performance between highly variable genes (HVGs) and non-highly variable genes (non-HVGs). For each gene, we first constructed predictive models of scRNA-seq via scATAC-seq using established methods. For a given gene, a higher prediction performance of the model indicated that the variability in expression of this gene between cells can be largely explained by chromatin accessibility. A significantly better prediction performance for HVGs than for non-HVGs means that the expression of HVGs is more strongly related to chromatin accessibility than that of non-HVGs. In other words, HVGs are subject to stricter epigenetic regulation than non-HVGs are. This provides evidence for the epigenetic regulation of scEV and underscores the role of epigenetic factors in determining scEV. As a result, we obtain reliable measurements of prediction performance and increase the robustness of our results.

We applied our pipeline to paired scATAC-seq and scRNA-seq datasets obtained from hematopoietic stem and progenitor cells (HSPCs). We found that the overall prediction performance of HVGs was indeed better than that of non-HVGs, which supports the epigenetic regulation of scEV. The results from further in-depth analysis provide additional evidence for the effects of chromatin accessibility on scEV. First, there was a significant correlation between the prediction performance and the level of scEV for different genes, indicating that chromatin accessibility is not independent of scEV. Second, there was a significant overlap between the well-predicted genes and HVGs, suggesting that chromatin accessibility plays a crucial role in the regulatory mechanism of HVGs. Therefore, we suggest that the variable expression of HVGs between cells is due to the varying levels of chromatin accessibility among these cells, and the regulation of HVGs can be inferred by analyzing the enriched transcription factor-binding sites in the scATAC-seq data. Finally, we validated the ability of scATAC-seq to reveal genes that indicate cell function through enrichment analysis of the well-predicted genes. In conclusion, our results provide single-cell evidence for the association between chromatin accessibility and scEV, highlighting the deterministic role of epigenetic mechanisms in modulating the stochasticity of gene expression. Our results also suggest a new approach for studying the mechanism of gene regulation by investigating how chromatin accessibility affects scEV.

## 2. Methods

There are four main steps in our testing pipeline, as shown in **Figure 1**: (1) collecting and preprocessing scATAC-seq and scRNA-seq datasets, (2) building predictive models on scRNA-seq data by scATAC-seq data and assessing the prediction performance of models for each gene, (3) calculating the level of scEV for each gene and assigning genes to the HVG or non-HVG groups, and (4) comparing the prediction performance of each model when applied to HVGs and non-HVGs.

**Figure 1.**
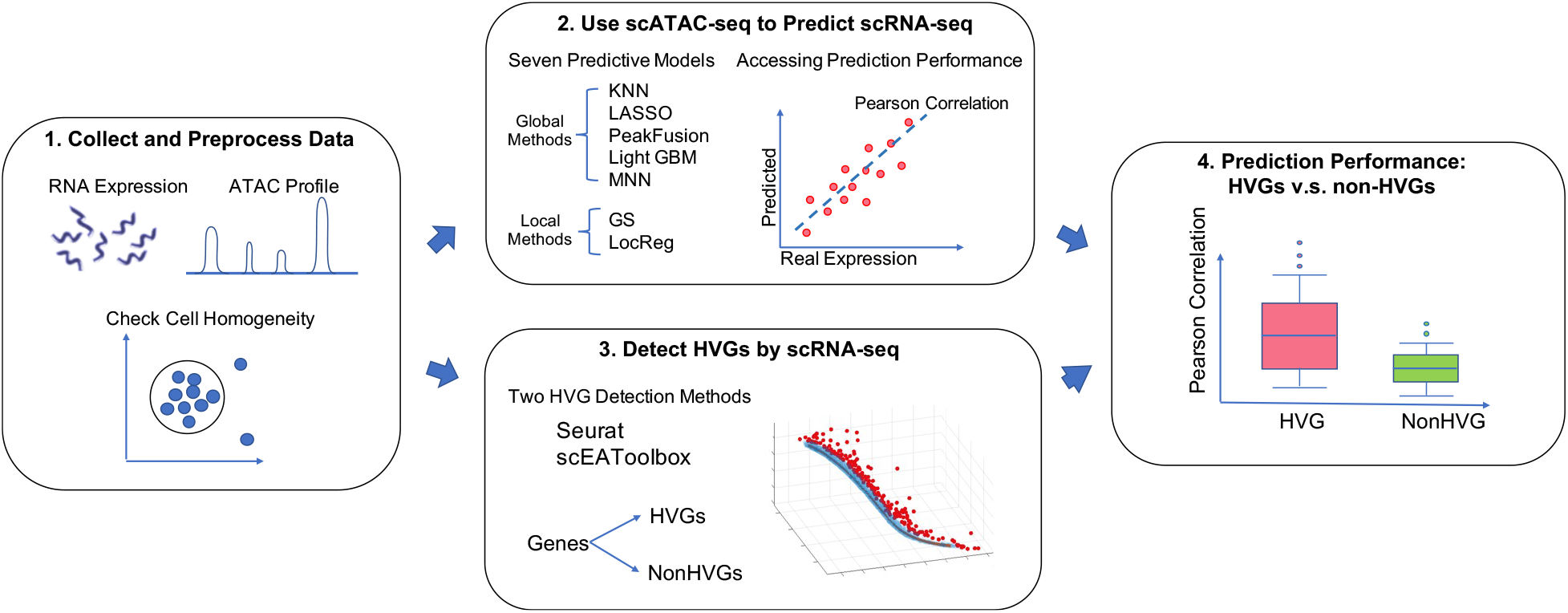
Overall pipeline for testing the relationship between chromatin accessibility and scEV.

### 2.1 Data collection and preprocessing

We used the paired scRNA-seq and scATAC-seq datasets provided by the Kaggle Open Problems in Single-Cell Analysis^1^. These datasets were derived from peripheral CD34+ hematopoietic stem and progenitor cells (HSPCs). To ensure cell homogeneity in our study, we focused on data from a single donor (donor #1) and a single time point (day 2). Starting with the scATAC-seq data, we removed cells that contained fewer than 1,000 scATAC-seq peaks. For the scRNA-seq data, we removed cells with fewer than 200 expressed genes and genes that were expressed in fewer than 200 cells or had a mean expression lower than 0.5. We also filtered out peaks and genes from the X, Y, and mitochondrial chromosomes, as well as those located within blacklisted genomic regions^2^[15]. Ultimately, we retained 6,871 cells, 206,611 peaks, and 9,657 genes for our analysis. The scATAC-seq peak counts were subsequently transformed using TF-IDF [16]. The scRNA-seq gene expression counts were sequentially library-size normalized and log1p transformed.

#### Check of Cell Homogeneity

To select homogeneous cells, we first extracted the first 50 principal components of the scATAC-seq data. Then, we utilized the UMAP algorithm [17] to obtain a two-dimensional representation of the cells. The scatter plot of cells in Figure 2 shows that the cells do not exhibit a homogeneous structure. Following a similar approach as Osorio et al. [5], we manually picked one cell from the center of the largest cluster as the core cell. We expanded the core by selecting the cells that were closest to the core cell until we had a total of 3,000 homogeneous cells (including the core cell itself). These selected cells were then used for further analysis.

**Figure 2.**
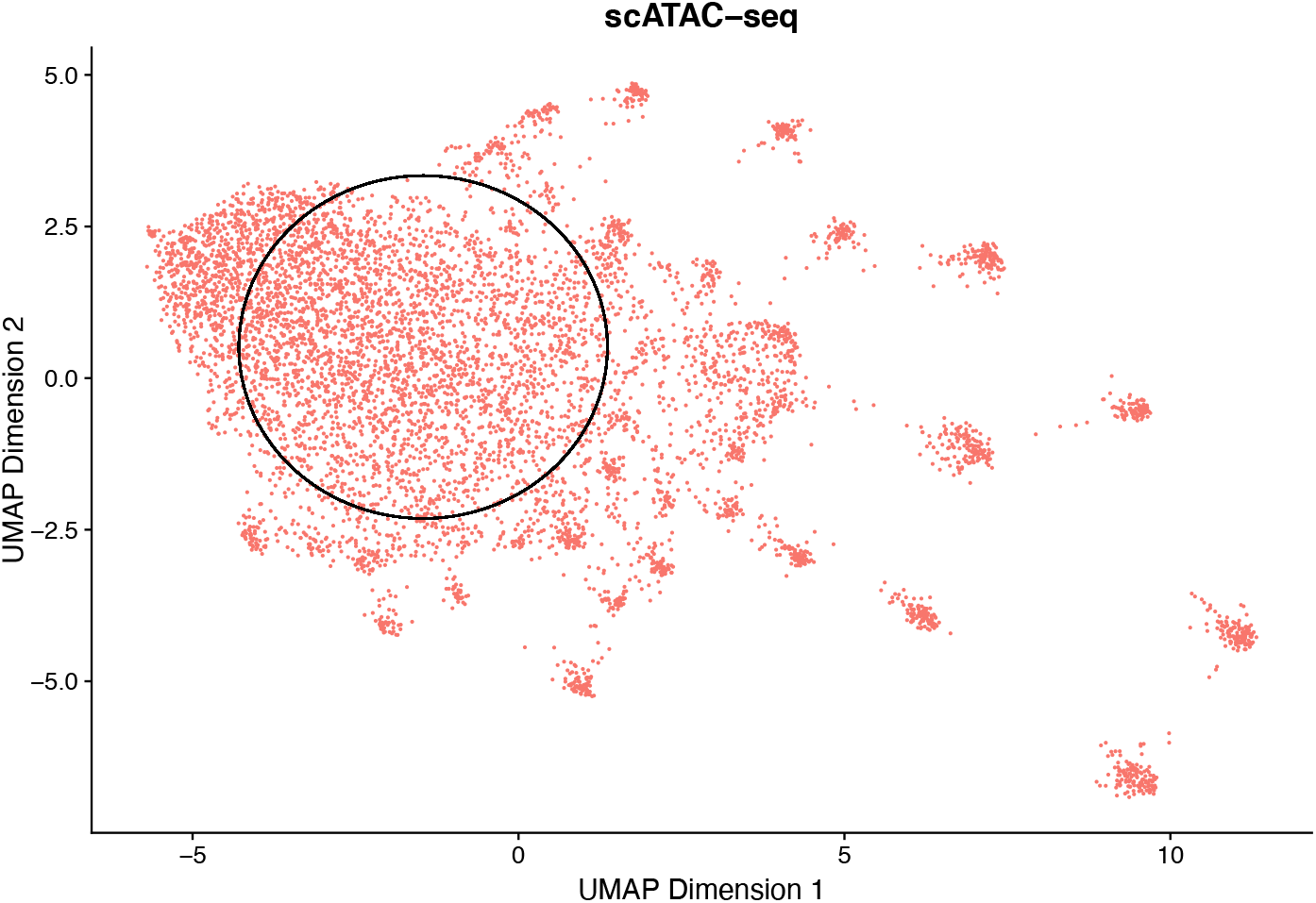
A homogeneous population of 3,000 cells was selected for analysis. UMAP was used to produce a 2-dimensional representation of the first 50 principal components of the scATAC-seq data. A core cell was chosen, and the 3,000 cells nearest to the core cell (including itself) were selected as a group of homogeneous cells. The selected cells are surrounded by the black circle.

### 2.2 Methods for predicting scRNA-seq gene expression with scATAC-seq peaks

Let *N, p*, and *q* be the number of cells, peaks, and genes, respectively, in the study. ***X*** ∈ ℝ^*N*×*p*^ and ***Y*** ∈ ℝ^*N*×*q*^ denote the scATAC-seq data matrix and the scRNA-seq data matrix, respectively. The *j*-th column of ***Y, y***_*j*_ ∈ ℝ^*N*^, includes the expression levels of the *j*-th gene in all cells. For each gene, we focused on building a predictive model on ***y***_*j*_ with ***X***.

Despite the availability of diverse predictive models, the use of scATAC-seq data for predicting scRNA-seq data remains in its early stages. To identify the most effective and reliable models, we explored seven different types of models that cover a wide range of established methods. These models can be classified into two categories: global and local methods. Global methods use all peaks as informative features for predicting gene expression, while local methods concentrate on peaks near the target gene on the same chromosome.

#### 2.2.1 Global methods

Five global methods were included and used in our analysis. We briefly describe models of these methods as follows.

1. K Nearest Neighbors (KNN): The KNN algorithm is a commonly used benchmarking model for scRNA data prediction [18]. For each cell, KNN first finds its *k* most similar cells according to scATAC-seq data; these cells are called its neighbors. Then, KNN predicts the expression level of a gene at this cell by averaging the expression levels of its neighbors. As the scATAC-seq data are extremely sparse, the similarity between two cells was calculated via their peak coexpression. *a*_1_, *a*_2_, *a*_3_ are the number of peaks where both cells have a nonzero count, only the first cell has a nonzero count, and only the second cell has a nonzero count, respectively. Then, we used *a*_1_/(*a*_1_ + *a*_2_ + *a*_3_) to measure the similarity between two cells. We selected *k* = 10 in the analysis for KNN.
2. Lasso [19]: Lasso is a popular linear regression model used to predict the expression levels of genes. It can identify important predictors that are most helpful for predicting the response variable. Specifically, Lasso minimizes the objective function of a squared loss term and a penalty term as follows:

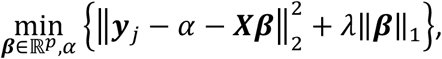

where *α* and ***β*** are the intercept and the coefficient vector of regression, respectively, and *λ* is a penalty parameter to control the sparsity of ***β***. After ***β*** is estimated, the peaks corresponding to the nonzero entries of ***β*** are those selected by Lasso that potentially affect the expression levels of the gene. We selected *λ* using the Bayesian information criterion (BIC) with the additional requirement that at least five peaks were selected by Lasso to avoid a trivial model.
3. Peak-Fusion-Based Model (PeakFusion) [20]: PeakFusion is a technique for addressing the high sparsity of scATAC-seq data, as more than 90% of the entries in the scATAC-seq matrix are zero. PeakFusion reduces the percentage of zeros in scATAC-seq data by summing the peaks within every 100 kb in chromosomes and forming an aggregated new predictor. The constructed new predictors were subsequently used to construct Lasso models to predict the expression levels of genes.
4. LightGBM [21]. LightGBM is a nonlinear machine learning method based on a gradient boosting decision tree. Compared with other boosting methods, LightGBM has a faster training speed and lower memory consumption due to the use of histogram optimization, the depth-first split strategy, the gradient-based one-sided sampling strategy, and the exclusive feature bundling strategy. We used the *sklearn* Python package (v.1.3.1) to construct LightGBM and set the number of trees to 10 while the other tuning parameters kept default.
5. Multilayer neural network (MNN). Neural networks have shown their power in numerical prediction problems. For each gene, we constructed a multilayer neural network comprising two hidden layers with the structure *p*-128-64-1. For each hidden layer, we added a dropout layer to prevent overfitting, for which the dropout rate = 0.4 [22]. The ReLU function was selected as the activation function, and the loss function was the mean square loss between the predicted expression level and the true expression level in the training dataset. Given that the number of peaks is very large, we also used the peak fusion technique described above to reduce the dimensionality of the input data and the computing time. All the input predictors were also normalized to [0,1] before training the model. This model is implemented via the *Keras* Python package (v.2.8.0).

#### 2.2.2 Local methods

Many existing studies [23, 10] suggest that the expression level of a gene is significantly influenced by nearby peaks on the same chromosome. As a result, we included two local prediction methods in our analysis.

1. Gene score (GS): GS is an unsupervised method introduced in ArchR [9]. In GS, the distance between the start of the target gene and the start of the peak is calculated. This distance is then converted to a distance weight by

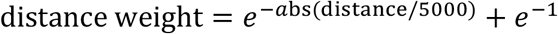 Next, the distance weight for the target gene is multiplied by the inverse of the gene size to account for differences in length among genes. The resulting value is then scaled from 1 to 5. Finally, the gene score for a gene in a cell is calculated by taking the elementwise product of the transformed distance weight and the peak. We calculated gene scores for all cells and treated them as predictions of gene expression levels.
2. Location information-based regression (LocReg): LocReg uses the nearby peaks of a gene as predictors to construct a Lasso model for predicting gene expression levels. In practice, we used the *biomaRt* R package (v.2.54.1) to obtain the location information of each gene. We considered only peaks within 1.2 Mbp of the gene for both local methods.

#### 2.2.3 Performance evaluation and selection of well-predicted genes

To evaluate the performance of the models in predicting gene expression, we randomly divided cells into a 70% training dataset and a 30% test dataset. The predictive models mentioned earlier were subsequently constructed using only the cells in the training dataset for each gene. These models were subsequently employed to predict the gene expression of cells in the test dataset. Pearson correlation between the actual and predicted gene expression levels in the test dataset was computed to evaluate the performance of each model. To ensure a reliable and robust evaluation, we repeated this random training-test sample splitting five times and obtained an average Pearson correlation as the metric for model evaluation. A higher average Pearson correlation coefficient for a given gene indicates that its expression level can be more accurately predicted using the scATAC-seq data.

Furthermore, for each prediction model, we ranked the genes based on their average Pearson correlation coefficient. The genes with the highest *K* Pearson correlation values were identified as well-predicted genes in our analysis, as their expression levels are primarily controlled by single-cell chromatin accessibility.

### 2.3 Measurement of scEV and identification of HVGs

Two established methods were chosen to measure the scEV levels of genes and identify HVGs from the scRNA-seq data. Both methods have been widely used and have shown effectiveness in previous studies.

1. Vst method in Seurat [24-27]: Vst uses locally estimated scatterplot smoothing to fit the relationship between the log of the variance and the log of the mean of gene expression levels. With the fitted variance and observed mean, the original data was normalized, and the new variance in each gene in the normalized data was considered a metric of scEV. A higher value indicates greater confidence in a gene being classified as an HVG.
2. Splinefit method in scGEAToolbox [28-30]: In addition to the mean and variance in gene expression levels, the splinefit method also considers the dropout rate of each gene and performs a spline fitting of a 3D curve using the three dimensions of dropout rate, log of the mean expression level, and log of the variance in the expression levels of all genes. After fitting the 3D curve, the distance from each gene to the curve was calculated and used as a metric of scEV. Genes that are farthest from the curve were identified as HVGs.

We applied both Seurat and scGEAToolbox to the scRNA-seq dataset. A systematic comparison of the results of the two methods is presented in **Figure 3a,b**. The Pearson correlation coefficient between the two different methods for determining the scEV levels of all the genes was as high as 0.79 as shown in **Figure 3c**. For each method, we ordered genes according to their scEV level in descending order. The top *K* genes were identified as HVGs, and the other genes were denoted as non-HVGs. When *K* = 200, the number of overlapping HVGs detected by both methods was as high as 177 (88.5%, **Figure 3d**). These results confirm the consistency of the scEV and HVG measurements obtained by the two methods.

**Figure 3.**
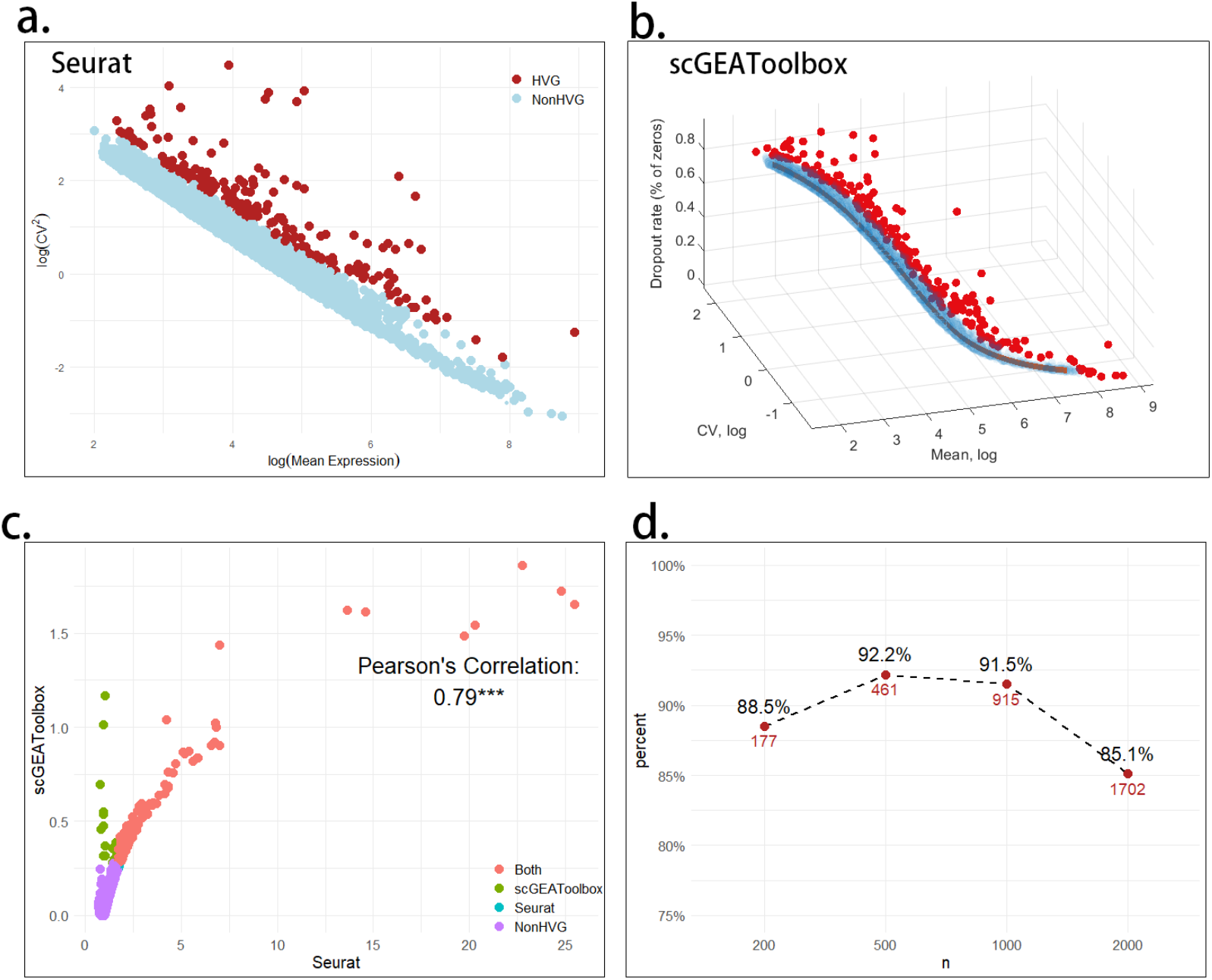
Results of two HVG detection methods. (a) The 2D scatterplot of genes with log(Mean Expression) as the x axis and log(CV^2^) as the y axis. Top *K* = 200 HVGs with relatively high log(CV^2^) corresponding to log(Mean Expression) are identified as HVG by Vst method in Seurat with red markers. (b) Splinefit method in scGEAToolbox draws the 3D scatterplot of genes in (log(Mean Expression), log(CV), Dropout rate) and fits a 3D curve. The *K* = 200 genes farthest from the curve are marked as the HVGs with red markers. (c) The scatterplot of the scEV levels of genes for two HVG detection methods. The Pearson correlation is 0.79 with p value <0.001. 177 genes simultaneously denoted as HVGs by both methods are colored in red. (d) The number and ratio of overlapped genes between two methods when selecting different numbers of HVGs (*K* = 200, 500, 1000, 2000).

### 2.4 Comparison of prediction performance when models are applied to HVGs and non-HVGs

To investigate the effects of chromatin accessibility on scEV, we compared the prediction results and scEV levels using three different tests. First, as discussed in the Introduction, we tested whether the overall prediction performance of HVGs was significantly greater than that of non-HVGs. Second, we calculated the Spearman correlation coefficient between the prediction performance and the scEV count and tested the significance of this correlation. Third, we identified K well-predicted genes and K HVGs and examined their overlap. We used Fisher’s exact test to determine the independence between the well-predicted genes and HVGs. Finally, we conducted a gene functional enrichment analysis of the well-predicted genes using enrichR [31] through its web interface to explore whether they are associated with cell-specific functions.

## 3. Results

### 3.1 Comparing the prediction performance of different predictive models

The predictive capabilities of all the genes across the seven different models are shown in **Figure 4**, which includes a boxplot illustrating the Pearson correlation coefficients of the genes (**Figure 4a**). Among the seven models, MNN demonstrated the best overall prediction performance across all genes, with a maximum correlation coefficient of approximately 0.7. In contrast, GS performed the worst, possibly because it was the only unsupervised method in the dataset.

**Figure 4.**
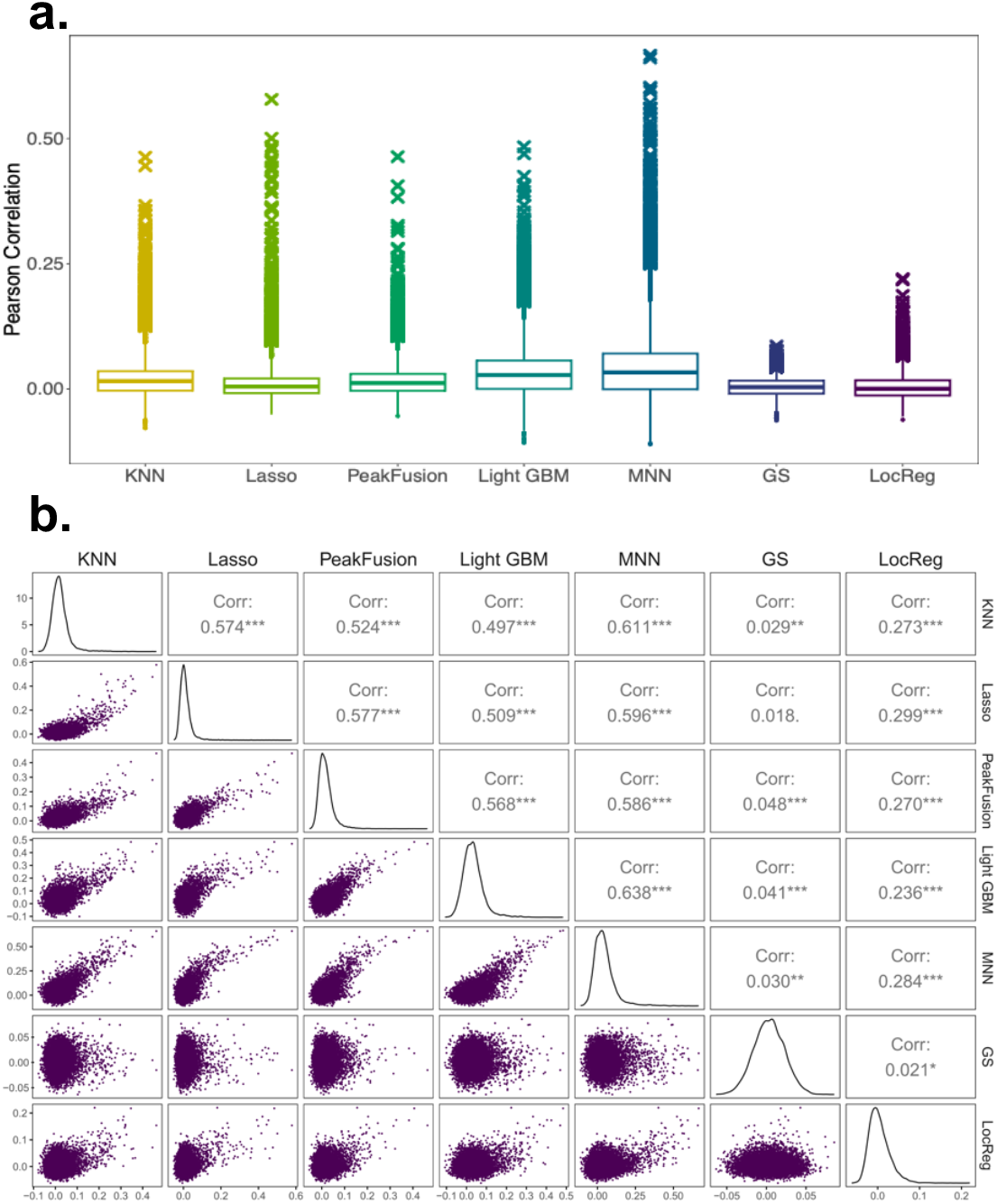
Results from seven prediction methods applied to scATAC-seq and scRNA-seq data. All the results represent the average results of five training-test sample splits. (**a**) Boxplot depicting Pearson correlations between true and estimated scRNA-seq levels for all genes across seven predictive models. The upper outliers in each boxplot signify genes that were well predicted with high Pearson correlations. In each model, the top 200 genes with the highest Pearson correlation coefficients are marked with a cross. (**b**) Scatter plots illustrating Pearson correlation for each pair of the seven models, with each point representing a gene.

Importantly, compared to local methods, global methods consistently outperform local methods, indicating a superior overall predictive capacity. These findings suggest that certain peaks, even those located at a distance from the target genes, control the expression levels of these genes. Therefore, these distant peaks should not be disregarded and should be included in predictive models.

The distributions of Pearson correlation coefficients on genes obtained using the seven models exhibit several common characteristics. Notably, the medians of Pearson correlation coefficients consistently hover approximately 0, indicating a general trend. However, some genes had relatively high Pearson correlation coefficients and were identified as outliers in the boxplot. This observation highlights that scATAC-seq data can accurately predict only a limited subset of genes, which we refer to as well-predicted genes in our analysis.

To evaluate the reliability of the prediction results, we calculated Pearson correlations between the prediction performances of each pair of models, as illustrated in **Figure 4b**. With the exception of pairs involving GS, all combinations of models demonstrated a statistically significant positive Pearson correlation (p value < 0.001), indicating a consistent predictive relationship between the scRNA-seq data and the scATAC-seq data across these models. Notably, MNN and LightGBM had the highest Pearson correlation (0.638), underscoring their particularly strong and aligned predictive capabilities.

### 3.2 Comparing prediction performance between HVGs and non-HVGs

Next, we compared the prediction performance of each predictive model when applied to HVGs and non-HVGs. We first identified *K* = 200 HVGs using either Seurat or scGEAToolbox. Boxplots were subsequently generated for each predictive model to show the average Pearson correlation for HVGs and non-HVGs (**Figure 5a**). The Wilcoxon test was used to compare the medians of the average Pearson correlations between the HVGs and non-HVGs. The overall prediction performance of HVGs is significantly greater than that of non-HVGs in all the models except for GS. The MNN model achieved the highest median Pearson correlations (0.26 and 0.25 for Seurat and scGEAToolbox, respectively) for HVGs. In contrast, the median Pearson correlation for non-HVGs in MNN was approximately 0. These results showed that within homogeneous cells, the scATAC data could provide much better predictions of the expression levels of HVGs than non-HVGs, suggesting that chromatin accessibility is a determining factor for scEV.

**Figure 5.**
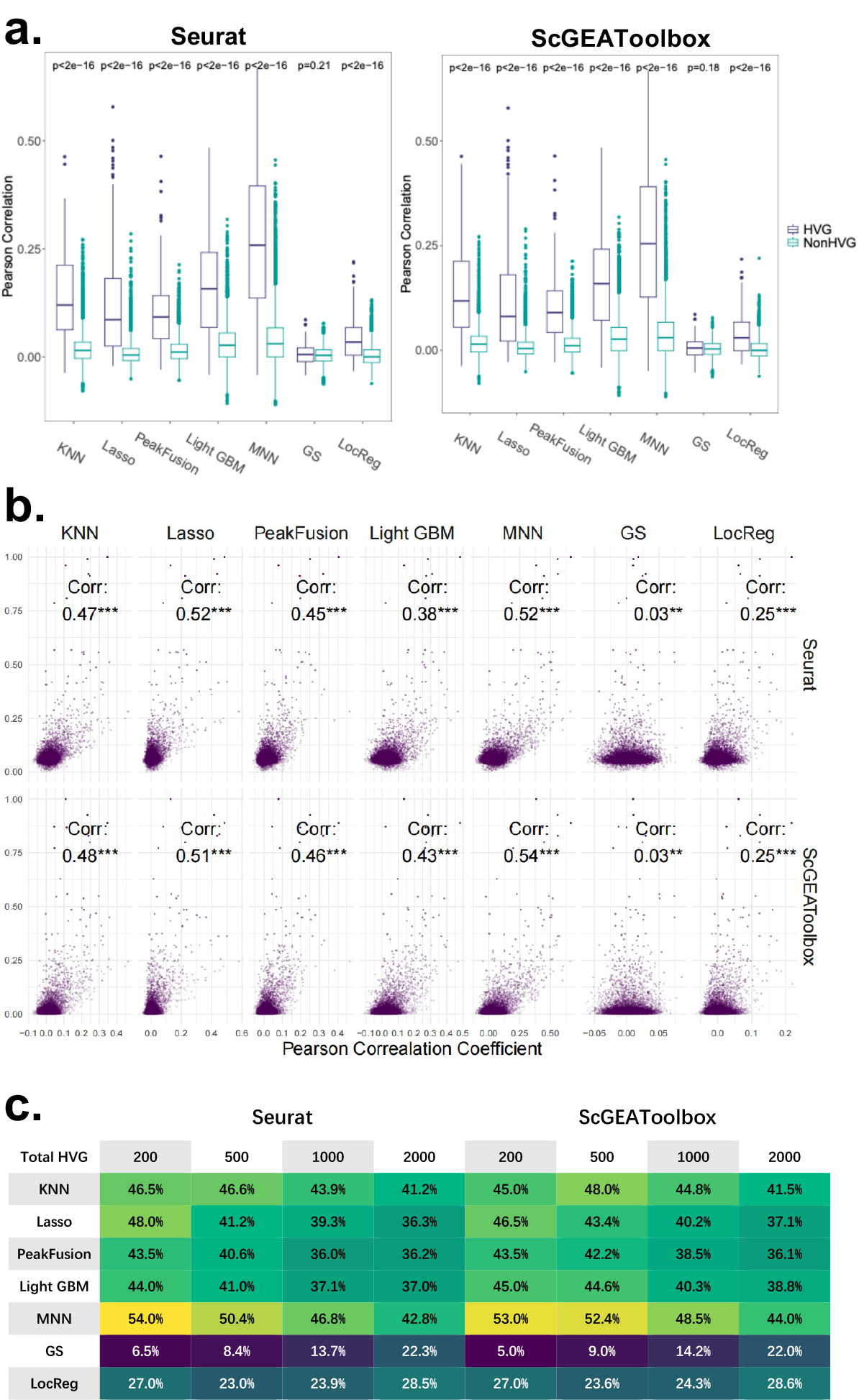
Prediction performance for HVGs across diverse models. (**a**) Identification of 200 HVGs using each HVG detection method, followed by boxplots illustrating the Pearson correlation for HVGs and non-HVGs within each predictive model. The Wilcoxon test was used to compare the medians of the two gene sets. (**b**) Scatterplots presenting genes with the x-axis indicating Pearson correlation and the y-axis representing scEV levels. Each column corresponds to a predictive model, and each row corresponds to an HVG detection method. Spearman correlation was calculated and tested for each scatterplot. (**c**) Evaluation of the overlap between the *K* well-predicted genes and the *K* detected HVGs for each pair of predictive models and HVG detection methods (*K* = 200, 500, 1000, 2000). Fisher’s exact test was used to assess the independence between well-predicted genes and HVGs for each method pair. All pairs demonstrated significant nonindependence, with a p value < 0.001, except for (GS, Seurat) and (GS, scGEAToolbox).

To further determine the relationship between chromatin accessibility and scEV, we systematically investigated the overall correlation between the average Pearson correlation coefficient and the scEV level for each gene according to each combination of predictive models and HVG detection methods. **Figure 5b** shows the scatterplot of genes in these two dimensions. A significant positive Spearman correlation was observed for all the combinations, except for those involving the GS model. The higher the predicted correlation coefficient is, the more likely a gene is to be considered an HVG. Notably, the Lasso and MNN methods achieved the highest Spearman correlations, surpassing 0.5.

Additionally, we examined the ability of chromatin accessibility to determine HVGs. For each predictive model, we selected the top *K* well-predicted genes with the best prediction performance as straightforward predictions for the top *K* HVGs. **Figure 5c** shows the overlap between the well-predicted genes and the detected HVGs for *K* = 200, 500, 1,000, and 2,000. Among all predictive models, the well-predicted genes by the MNN model exhibited the highest overlap with HVGs, accounting for 54% for Seurat and 53% for scGEAToolbox when *K* = 200. Therefore, utilizing well-predicted genes directly can recover more than 50% of HVGs. The results of Fisher’s exact tests indicate that well-predicted genes and HVGs, provided by all combinations of predictive models and HVG detection methods, except those involving the GS model, are not independent, providing evidence for the effect of chromatin accessibility on scEV control.

### 3.3 Well-predicted genes reveal cell type-specific functions

Given that the well-predicted genes significantly overlapped with the detected HVGs, we investigated whether these well-predicted genes are associated with cell type-specific functions. Among the seven predictive models, we selected Lasso and MNN as representative global methods and LocReg as a representative local method. We selected 200 well-predicted genes predicted by the Lasso, MNN, and LocReg models and performed gene enrichment analysis.

The top ten pathways associated with genes whose expression was most significantly altered are depicted in **Figure 6**. Many of these pathways are related to the functions of HSPCs. For example, *Hemostasis* is ranked first in both MNN and Lasso and third in Local. Hemostasis is the important function of HSPCs because it plays a critical role in maintaining the balance between bleeding and clotting, which is essential for overall vascular health [32]. HSPCs are responsible for replenishing blood cell populations, including platelets, which aid in hemostasis [33]. Identification of the hemostasis gene set in HSPCs helps to reveal the regulatory mechanisms that govern the delicate balance of coagulation and anticoagulation processes within the hematopoietic system, and to understand the complex molecular pathways that govern the fate and function of HSPCs. *Neutrophil degranulation*, highly ranked by all the methods, is also essential due to its role in the immune response and hematopoiesis. HSPCs give rise to various blood cell lineages, including neutrophils [34]. Neutrophils degranulation is a fundamental process in which these immune cells release granules containing antimicrobial proteins, enzymes, and reactive oxygen species [35]. Identification of genes regulating neutrophil degranulation in HSPCs provides insights into the complex mechanisms governing immune cell development and function in the host defense system.

**Figure 6.**
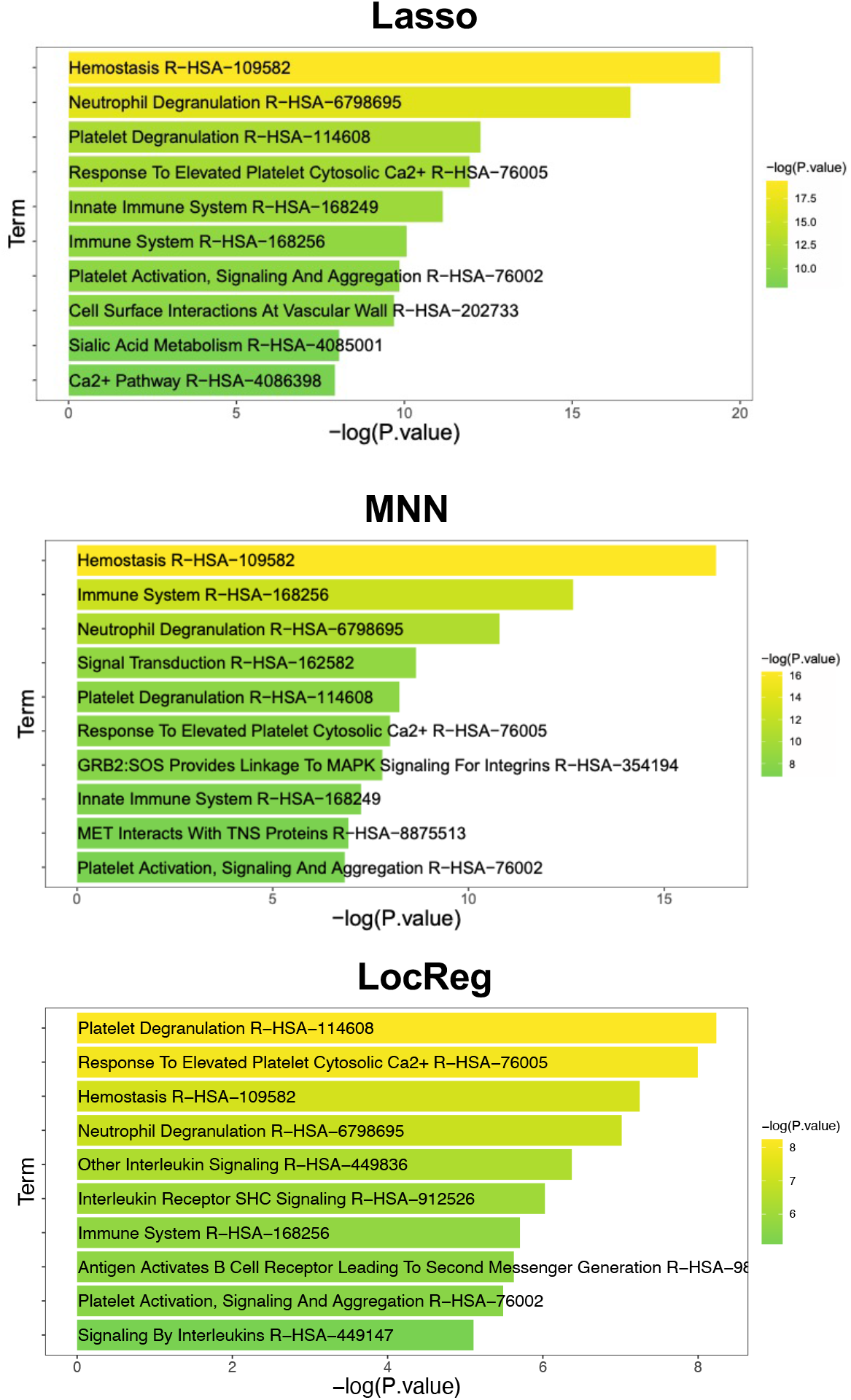
Lists of the top 10 significant pathways of the top 200 well-predicted genes according to the Lasso, MNN, and LocReg models.

### 3.4 Sensitivity analysis

To demonstrate the robustness of our analysis results, we assessed whether the prediction performance in this study was influenced by the mean expression level of the genes. For each of the 200 HVGs detected by Seurat, we selected the non-HVG with the most similar mean expression level for comparison. **Figure 7** displays the scatterplot of the average Pearson correlation coefficient between each HVG and its corresponding non-HVG. The plot indicates that HVGs generally exhibit significantly better prediction performance than their non-HVG counterparts. Therefore, our findings were not influenced by the mean expression level.

**Figure 7.**
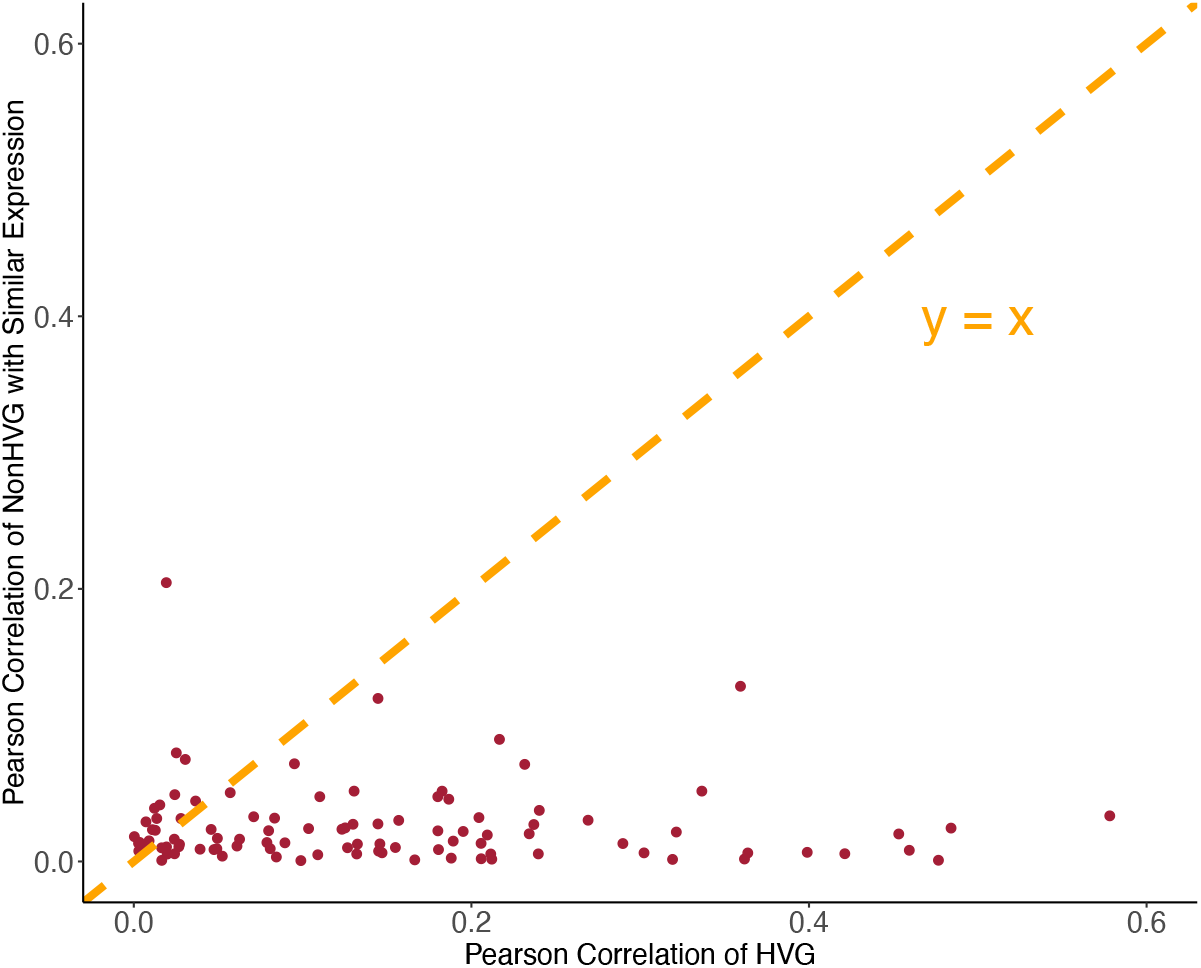
Prediction performance of HVGs and non-HVGs with controlled mean expression level. For each of *K* = 200 HVGs by Seurat, the non-HVG with the closest mean expression level is used as a compared gene with mean expression level controlled. The scatterplot of the average Pearson correlation of each HVG and its compared non-HVG shows that HVG have much better prediction performance than its compared non-HVG.

## 4 Discussion

In this study, we presented a new testing pipeline to study the relationship between scATAC-seq data and scEV. We specifically focused on homogeneous cells and found that genes whose expression levels could be accurately predicted by scATAC-seq were more likely to be HVGs. These findings support the notion of epigenetic regulation of scEV. Additionally, our results suggest that chromatin accessibility may play a role in gene regulation by influencing the cell-to-cell variability in gene expression. This provides valuable insights for studying gene function through analyzing scATAC-seq data. We also applied our testing pipeline to several other paired scATAC-seq and scRNA-seq datasets. While detailed results are omitted for brevity, the additional datasets we analyzed reinforced the notion that scATAC-seq possesses superior predictive power for HVGs compared to non-HVGs across diverse cellular contexts. Moreover, genes demonstrating high prediction accuracy consistently mapped to cell type-specific functions.

In summary, the emergence of powerful single-cell techniques has paved the way for a deeper understanding of the mechanisms driving scEV. Our findings offer compelling evidence that cell-to-cell variations in chromatin accessibility are a major driver of gene expression noise. This not only confirms the epigenetic regulation of noise but also opens exciting avenues for exploring gene regulation mechanisms at the single-cell level.

## Data availability

All the data used in this article are available from public sources, as detailed above in Section 2.1.

## Footnotes

1 https://www.kaggle.com/competitions/open-problems-multimodal/data.

2 https://sites.google.com/site/anshulkundaje/projects/blacklists.

## Reference

[1] Wada, T., Hironaka, K. I., Wataya, M., Fujii, M., Eto, M., Uda, S., … & Kuroda, S. (2020). Single-cell information analysis reveals that skeletal muscles incorporate cell-to-cell variability as information not noise. Cell Reports, 32(9).

[2] Snijder, B., & Pelkmans, L. (2011). Origins of regulated cell-to-cell variability. Nature Reviews Molecular Cell Biology, 12(2), 119–125.

[3] Zheng, H., Vijg, J., Fard, A. T., & Mar, J. C. (2023). Measuring cell-to-cell expression variability in single-cell RNA-sequencing data: a comparative analysis and applications to B cell aging. Genome biology, 24(1), 238.

[4] Dueck, H., Eberwine, J., & Kim, J. (2016). Variation is function: Are single cell differences functionally important? Testing the hypothesis that single cell variation is required for aggregate function. BioEssays, 38(2), 172–180.

[5] Osorio, D., Yu, X., Zhong, Y., Li, G., Serpedin, E., Huang, J. Z., & Cai, J. J. (2020). Single-cell expression variability implies cell function. Cells, 9(1), 14.

[6] Wiggins, G. A., Black, M. A., Dunbier, A., Morley-Bunker, A. E., kConFab Investigators, Pearson, J. F., & Walker, L. C. (2021). Increased gene expression variability in BRCA1-associated and basal-like breast tumours. Breast Cancer Research and Treatment, 189, 363–375.

[7] Kartha, V. K., Duarte, F. M., Hu, Y., Ma, S., Chew, J. G., Lareau, C. A., … & Buenrostro, J. D. (2022). Functional inference of gene regulation using single-cell multi-omics. Cell Genomics, 2(9).

[8] Stuart, T., Srivastava, A., Madad, S., Lareau, C. A., & Satija, R. (2021). Single-cell chromatin state analysis with Signac. Nature Methods, 18(11), 1333–1341.

[9] Granja, J. M., Corces, M. R., Pierce, S. E., Bagdatli, S. T., Choudhry, H., Chang, H. Y., & Greenleaf, W. J. (2021). ArchR is a scalable software package for integrative single-cell chromatin accessibility analysis. Nature Genetics, 53(3), 403–411.

[10] Mitra, S., Malik, R., Wong, W., Rahman, A., Hartemink, A. J., Pritykin, Y., … & Leslie, C. S. (2023). Single-cell multiome regression models identify functional and disease-associated enhancers and enable chromatin potential analysis. bioRxiv, 2023–06.

[11] Saelens, W., Pushkarev, O., & Deplancke, B. (2023). ChromatinHD connects single-cell DNA accessibility and conformation to gene expression through scale-adaptive machine learning. bioRxiv, 2023–07.

[12] Li, B., & You, L. (2013). Predictive power of cell-to-cell variability. Quantitative Biology, 1(2), 131–139.

[13] Eling, N., Morgan, M. D., & Marioni, J. C. (2019). Challenges in measuring and understanding biological noise. Nature Reviews Genetics, 20(9), 536–548.

[14] Buenrostro, J. D., Wu, B., Litzenburger, U. M., Ruff, D., Gonzales, M. L., Snyder, M. P., … & Greenleaf, W. J. (2015). Single-cell chromatin accessibility reveals principles of regulatory variation. Nature, 523(7561), 486–490.

[15] Carroll, T. S., Liang, Z., Salama, R., Stark, R., & de Santiago, I. (2014). Impact of artifact removal on ChIP quality metrics in ChIP-seq and ChIP-exo data. Frontiers in Genetics, 5, 75.

[16] Chowdhury, G. G. (2010). Introduction to modern information retrieval. Facet publishing.

[17] McInnes, L., Healy, J., Saul, N., & Großberger, L. (2018). UMAP: Uniform Manifold Approximation and Projection. Journal of Open Source Software, 3(29), 861.

[18] Wu, K. E., Yost, K. E., Chang, H. Y., & Zou, J. (2021). BABEL enables cross-modality translation between multiomic profiles at single-cell resolution. Proceedings of the National Academy of Sciences, 118(15), e2023070118.

[19] Tibshirani, R. (1996). Regression shrinkage and selection via the lasso. Journal of the Royal Statistical Society Series B: Statistical Methodology, 58(1), 267–288.

[20] Cao, J., Cusanovich, D. A., Ramani, V., Aghamirzaie, D., Pliner, H. A., Hill, A. J., … & Shendure, J. (2018). Joint profiling of chromatin accessibility and gene expression in thousands of single cells. Science, 361(6409), 1380–1385.

[21] Ke, G., Meng, Q., Finley, T., Wang, T., Chen, W., Ma, W., … & Liu, T. Y. (2017). Lightgbm: A highly efficient gradient boosting decision tree. Advances in Neural Information Processing Systems, 30.

[22] Srivastava, N., Hinton, G., Krizhevsky, A., Sutskever, I., & Salakhutdinov, R. (2014). Dropout: a simple way to prevent neural networks from overfitting. The Journal of Machine Learning Research, 15(1), 1929–1958.

[23] Alanis-Lobato, G., Bartlett, T. E., Huang, Q., Simon, C. S., McCarthy, A., Elder, K., … & Niakan, K. K. (2024). MICA: a multi-omics method to predict gene regulatory networks in early human embryos. Life Science Alliance, 7(1).

[24] Hao, Y., Hao, S., Andersen-Nissen, E., Mauck, W. M., Zheng, S., Butler, A., … & Satija, R. (2021). Integrated analysis of multimodal single-cell data. Cell, 184(13), 3573–3587.

[25] Stuart, T., Butler, A., Hoffman, P., Hafemeister, C., Papalexi, E., Mauck, W. M., … & Satija, R. (2019). Comprehensive integration of single-cell data. Cell, 177(7), 1888–1902.

[26] Butler, A., Hoffman, P., Smibert, P., Papalexi, E., & Satija, R. (2018). Integrating single-cell transcriptomic data across different conditions, technologies, and species. Nature Biotechnology, 36(5), 411–420.

[27] Satija, R., Farrell, J. A., Gennert, D., Schier, A. F., & Regev, A. (2015). Spatial reconstruction of single-cell gene expression data. Nature Biotechnology, 33(5), 495–502.

[28] Cai, J. J. (2020). scGEAToolbox: a Matlab toolbox for single-cell RNA sequencing data analysis.

[29] Sheng, J., & Li, W. V. (2021). Selecting gene features for unsupervised analysis of single-cell gene expression data. Briefings in Bioinformatics, 22(6), bbab295.

[30] Gilis, Jeroen, Laura Perin, Milan Malfait, Koen Van den Berge, Alemu Assefa Takele, Bie Verbist, Davide Risso, and Lieven Clement. “Differential detection workflows for multi-sample single-cell RNA-seq data.” bioRxiv (2023): 2023–12.

[31] Kuleshov, M. V., Jones, M. R., Rouillard, A. D., Fernandez, N. F., Duan, Q., Wang, Z., … & Ma’ayan, A. (2016). Enrichr: a comprehensive gene set enrichment analysis web server 2016 update. Nucleic Acids Research, 44(W1), W90–W97.

[32] Nguyen, T. S., Lapidot, T., & Ruf, W. (2018). Extravascular coagulation in hematopoietic stem and progenitor cell regulation. Blood, The Journal of the American Society of Hematology, 132(2), 123–131.

[33] Ouseph, M. M., Huang, Y., Banerjee, M., Joshi, S., MacDonald, L., Zhong, Y., … & Wang, Q. J. (2015). Autophagy is induced upon platelet activation and is essential for hemostasis and thrombosis. Blood, The Journal of the American Society of Hematology, 126(10), 1224–1233.

[34] Iwasaki, H., & Akashi, K. (2007). Myeloid lineage commitment from the hematopoietic stem cell. Immunity, 26(6), 726–740.

[35] Malech, H. L., DeLeo, F. R., & Quinn, M. T. (2014). The role of neutrophils in the immune system: an overview. Neutrophil Methods and Protocols, 3–10.

